# Exploring connectivity of resting-state EEG between BCI-literate and -illiterate groups

**DOI:** 10.1101/2023.11.25.568691

**Authors:** Hanjin Park, Sung Chan Jun

**Affiliations:** AI Graduate School, Gwangju Institute of Science and Technology, Gwangju, South Korea; School of Electrical Engineering and Computer Science, Gwangju Institute of Science and Technology, Gwangju, South Korea

## Abstract

Although Motor Imagery-based Brain-Computer Interface (MI-BCI) holds significant potential, its practical application faces challenges attributable to the phenomenon known as BCI-illiteracy. BCI researchers have attempted to predict BCI-illiteracy to mitigate this issue. As connectivity’s significance in neuroscience has grown, BCI researchers have applied connectivity to predict BCI-illiteracy with resting-state data. However, connectivity metrics’ use and interpretation of the results can be challenging for several reasons. Firstly, there are various connectivity metrics, each with its own advantages and disadvantages based on their underlying hypotheses and perspectives. These pros and cons are shaped by several factors, increasing the complexity of their application and interpretation. Secondly, it is unclear whether they are as acceptable as their developers have claimed. Thirdly, it is not evident which factor may influence the estimation of connectivity and which metric is suitable for this research. Therefore, this study conducted an empirical test to provide BCI researchers with a better understanding of connectivity. We analyzed three large public datasets using three functional connectivity (FC) and three effective connectivity (EC) metrics. Additionally, the structural difference in the resting-state network between BCI-literate and illiterate groups was examined. Our analysis revealed that the appropriate frequency range to measure connectivity varies depending upon the metric used. The alpha range was found to be suitable for FC, while the alpha, alpha + theta, and beta ranges were found to be appropriate for EC. Further, the results of estimating connectivity varied depending upon the dataset and metric used. Although we observed that BCI-literacy had stronger connections between nodes, no other significant structural differences were found between the two groups. However, BCI-literacy’s resting-state network displayed higher network efficiency compared to BCI-illiteracy, regardless of the metrics and dataset used. Therefore, it seems reasonable to use resting-state connectivity to predict BCI-illiteracy. Our conclusion is that each metric has its own specific hypothesis and perspective to measure connectivity under certain conditions.

## 1 Introduction

Motor imagery-based brain-computer interface (MI-BCI) is a promising technology that allows communication with external devices by imagining body movements rather than performing them physically [1]. MI-BCI promises that individuals with physical disabilities can interact with their surroundings effectively. For example, a person with a physical disability would receive assistance from a wheelchair or robotic arm [2]. The brain signals for BCI input can be collected through electroencephalography (EEG). MI-BCI works by detecting unique brain signal patterns when different body movements are imagined [3]. For a successful MI-BCI, MI’s classification accuracy must surpass the criterion level [4]–[8]. However, it has been reported that approximately 15-30% of BCI users cannot achieve the accuracy rate necessary for successful MI-BCI [9], [10]. These users are defined as ‘BCI-illiterate’ or ‘BCI-inefficient’ [11].

BCI researchers have investigated various ways to identify BCI-illiteracy before they conduct resource-intensive BCI experiments. Researchers are particularly interested in the ‘Resting state’, where a subject does nothing with open or closed eyes, as sensorimotor rhythm (SMR) is associated with a resting state. SMR is suppressed during motor execution and MI [12]. Such a phenomenon is referred to as Event-Related Desynchronization (ERD), which is considered an essential property of MI [13]. Blankerts et al. reported that a lower power of SMR during a resting state contributes to BCI-illiteracy [12]. Ahn et al. found that the higher relative alpha power and lower relative theta power during the resting state may predict BCI-illiteracy [14]. Kwon et al. also suggested that BCI-illiteracy may be associated with the relative beta power [15]. According to Grosse-Wentrup et al., the level of gamma power during the resting state could have an effect on BCI performance by influencing SMR [16]. In addition, Zhang et al. discovered a positive correlation between BCI performance and the resting state using the spectral entropy [17]

The studies above have focused on specific channels, taking a ‘Functional specialization’ perspective [18]. However, the brain operates as a dynamic network, which cooperates dynamically with diverse brain regions to perform tasks or processes [19], [20]. From this perspective, the microscopic approach provides only a limited understanding of brain function. As numerous studies on resting-state brain functional networks have revealed previously unknown brain functions [21], [22], BCI researchers have taken into account the functional network to identify what contributes to BCI-illiteracy. Zhang et al. reported that BCI performance could be predicted by analyzing the resting-state EEG network’s efficiency [23]. Li et al. suggested that effective reconfiguration of the brain from resting state to MI task execution is related to BCI performance [24]. Lee et al. found that the strength of connectivity in the resting-state network may indicate BCI-illiteracy [25]. Leeuwis et al. revealed the feature of BCI-illiteracy by comparing the brain network of a resting state and that of an MI task[26]. Previous research has focused on whether resting-state functional brain networks can predict BCI performance. However, the resting state network has not been investigated thoroughly beyond examining the way that certain graph features are correlated with BCI performance. As a result, it remains unclear what similarities and differences exist in the structure of resting state networks between BCI-literacy and illiteracy. This is particularly disappointing given the growing significance of the default mode network (DMN), brain regions that activate during resting state, in neuroscience [27]–[29].

Analyzing the brain’s ‘connectivity’ is an important aspect of researching brain functional networks [22], and numerous methodologies have been developed to quantify and analyze connectivity to date. However, as Bastos et al. pointed out, because each connectivity metric has its own hypothesis, it has its own advantages and disadvantages [19]. As a result, even estimating connectivity with the same dataset may vary depending upon metrics. In addition, the experimental context and data characteristics influence the metrics. If a researcher fails to contemplate this, it could lead to an incorrect interpretation [19]. Thus, it is recommended to compare results from various connectivity metrics to ensure accuracy. Colclough et al. studied which connectivity metric produces the most dependable outcome on resting-state MEG data [30]. However, to the best of our knowledge, the BCI studies addressed above used two metrics at most and did not provide a comprehensive analysis of the results with respect to the metrics used.

Further, it has remained unclear which frequency band is crucial to identify BCI-illiteracy through connectivity analysis. Zhang et al. used a frequency range of 4-14 Hz to compute a connectivity matrix based upon the research that indicates that the theta and alpha bands are relevant to BCI-illiteracy [23]. However, this finding was drawn from analyzing band power, not connectivity. Thus, it is uncertain which band is most pertinent to BCI-illiteracy in connectivity and whether or not it is appropriate to combine the two bands. It is possible that the proper band varies depending upon the metric. Therefore, exploring each metric’s proper frequency range in a specific research context is worthwhile.

As mentioned earlier, the results of connectivity metrics may depend upon the dataset used, such that the result is valid only with a specific dataset. Hence, it is crucial to replicate the result using different datasets. However, to the best of our knowledge, replicating the results of estimating connectivity with other datasets is rarely conducted in MI-BCI research [23]–[26].

Our goal in this study is to offer a better understanding and perspective on resting-state EEG connectivity by investigating the degree to which the resting-state network predicts future BCI performance, and analyzing BCI-literacy and -illiteracy’s networks. We assumed that the two groups have distinct resting-state EEG based upon a previous study [14]. First, we compared the outcomes of three functional and three effective connectivity metrics commonly employed for estimation with a large dataset (N = 52) [31]. We believe that comparing metrics helps a researcher gain a practical understanding and intuition of connectivity. Second, we explored each metric’s appropriate frequency range to identify BCI-illiteracy. We expect that identifying the frequency range’s effect on estimating connectivity will expand our knowledge about connectivity. Third, we conducted an in-depth analysis of the resting-state brain network using diverse graph theory measures, and we focused in particular on identifying the structural difference between BCI-literacy and -illiteracy. Fourth, we replicated our study with two larger datasets (N = 54 x 2 sessions) with similar experimental settings to provide more comprehensive results [32]. This comparison enhanced not only our results’ robustness, but also allowed us to evaluate the dataset’s effect on our estimation of connectivity. Finally, we integrated all three datasets [31], [32] and investigated whether the findings above can be captured in the entire dataset.

## 2 Materials and Methods

### 2.1 EEG datasets

We used two publicly available EEG datasets of BCI experiments that focused on MI of the left and right hands. The primary dataset for this study was from Cho et al., and comprised 52 subjects (19 females, 24.8 ± 3.86 years) [31]. The dataset consisted of 100 or 120 offline MI trials for each hand, as well as 1 minute of resting state data before the experiment. EEG data were collected from 64 active electrodes based upon the international 10-10 system with a 512 Hz sampling rate.

The second dataset comprised 54 subjects (25 females, 24.8 ± 3.8 years) [32]. The experiments were performed over two sessions on different days. This dataset included 50 offline MI trials for each hand and 1 minute of resting state data before the experiment. EEG data were recorded using 62 electrodes based upon the International 10-20 system with a 1000 Hz sampling rate.

### 2.2 Evaluation of BCI Performance and Group Categorization

For this study, we followed the process and parameters established by the dataset’s authors closely to evaluate BCI performance and focused on analyzing the resting-state connectivity. In the primary dataset, all 64 channels were used. The MI EEG data were pre-processed by a common average reference (CAR). Then, we band-pass filtered between 8-30 Hz and extracted 0.5-2.5 seconds of MI data from a 3-second duration to evaluate performance with all channels. We used the common spatial pattern (CSP) to extract features for each class, and Fisher’s Linear discriminant analysis (LDA) as the classifier. For the evaluation, we executed 10-fold cross-validation with 7 training sets and 3 testing sets. A mean accuracy of 120 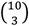 times estimation was used as a subject’s BCI performance [31].

For the second dataset, we used 1-3.5 seconds of MI EEG data of a 4-second duration, and 20 channels (FC1, FC2, FC3, FC4, FC5, FC6, Cz, C1, C2, C3, C4, C5, C6, CPz, CP1, CP2, CP3, CP4, CP5, and CP6), as the author did. In addition, we down-sampled the second dataset’s sampling rate to 500 to make it similar to that of the primary dataset [32]. The remaining steps for the evaluation were the same.

With the estimated BCI performance, we categorized subjects into two distinct groups: the ‘BCI-literate’ group and the ‘BCI-illiterate’ group. We defined a subject as BCI-illiteracy if their performance did not exceed the chance level. The chance level in binary classification varies greatly depending upon the number of trials [33], [34]. Hence, it was necessary to establish specific thresholds to determine BCI-illiteracy that correspond to the different number of trials. For the primary dataset, in which the number of trials is 100 or 120 for each class, the threshold is under 60% accuracy. The threshold for the second dataset is 62% accuracy, as the number of trials is 50 for each class. Thus, subjects with an accuracy below 60% in the primary dataset or below 62% in the second dataset were considered BCI-illiteracy. However, we used the same threshold to determine BCI-literacy, because it does not relate to defining the chance level criterion. We defined a subject whose accuracy exceeded 70% in both datasets as BCI-literate. This threshold is recognized as an appropriate threshold for BCI-literacy and indicates a significant improvement from the chance level [4]–[8].

### 2.3 Connectivity

The metric can be categorized based upon whether it provides directional (or causal) information or not [19]. Functional connectivity (FC) is an undirected metric that captures the statistical dependence between signals. FC captures only statistical interdependence between signals-of-interest. Cross-correlation, Coherence (COH), Imaginary Coherence (iCOH), and Phase Locking Value (PLV) are popular FC metrics. On the other hand, effective connectivity (EC) measures causal interaction between signals to assess information exchange. EC includes Granger Causality (GC), Transfer Entropy (TE), and Phase Slope Index (PSI). GC and TE provide bidirectional information by estimating each signal’s predictability with respect to rest signals [19], [35]–[37]. However, PSI measures only unidirectional interaction between signals [19], [38]. The connectivity estimation can occur in either the frequency or time domain, depending upon the specific metric used [19]. Frequency domain metrics, such as COH, iCOH, PLV, and PSI, are typically employed to measure connectivity based upon phase synchronization between signals [25,40–42]. These metrics reveal the way that signals interact and synchronize at different frequency bands. In contrast, time-domain metrics, like Cross-Correlation, GC, and TE, analyze the data’s temporal structure. These metrics examine the temporal relations or causal interactions between signals [19], [35]–[37]. GC and TE take a similar perspective, in which the causal relation is revealed through conditional probability to measure the causality between signals [35]–[37], [39]. GC can also be estimated in the frequency domain, and its application can be subdivided based upon theoretical underpinnings [19], [35], [36]. In this study, we employed COH, iCOH, PLV, PSI, Bivariate GC (BVGC), and Multivariate GC (MVGC). In addition, we considered GC only in the time domain to focus on comparing the BVGC and MVGC’s properties.

#### 2.3.1 The procedure to estimate connectivity metrics in the frequency domain

15 channels (F3, F4, FC3, FC4, Cz, C3, C4, C5, C6, CP3, CP4, P3, P4, O1, and O2) were selected to estimate both datasets’ resting state EEG data [23]. 60-second resting state EEG data were segmented into 2 seconds, as we concluded that this was the best epoch size to evaluate connectivity. However, the epoch size influenced the estimation result only slightly [30].

We preprocessed the data to analyze frequency using the Fieldtrip toolbox [40]. The data were filtered between 1-60 Hz (1-80 Hz for the gamma band) and re-referenced using a CAR. Multi-Taper Method Fast Fourier Transform (MTFFT) was executed with Discrete Prolate Spheroidal Sequences (DPSS) and 3 Hz smoothing. We calculated connectivity values between channels within theta (4-8 Hz), alpha (8-13 Hz), theta + alpha (4-13 Hz), beta (13-30 Hz), and gamma (30-70 Hz) range separately [14]. After evaluating a connectivity matrix with the values, we conducted block permutation bootstraps and False Discovery rate (FDR) with alpha = 0.05 to reduce spurious connectivity values. However, we did not perform the block permutation bootstraps for PSI because of the metric’s property [38]. After completing these steps, we obtained a single connectivity matrix for 2-second resting state data. Finally, we averaged 30 matrices to create a connectivity matrix for analysis. The procedure is depicted in Fig. 1.

**Fig. 1.**
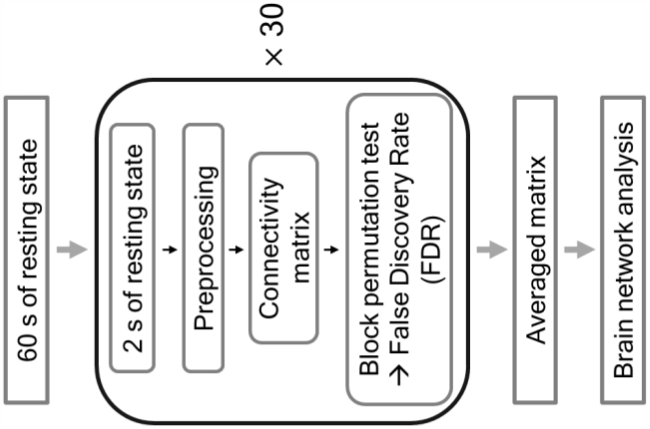
The procedure of connectivity estimation in the frequency domain.

**Coherence** COH is a metric used widely to quantify the synchronization between two signals. Mathematically, COH is a cross-correlation function in the frequency domain [19]. COH indicates how much one signal’s variance is explained by another or the converse. The value ranges between 0 and 1. 1 indicates that the two signals are synchronized perfectly, while 0 indicates that they are not synchronized at all.

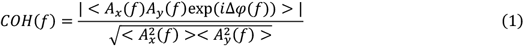

In this equation, *A*_*i*_ stands for the amplitude of signal *i*, and *f* signifies the frequency range of interest. The term Δ*ϕ*(*f*) is the phase difference between the two signals for frequency *f*. The numerator represents an average cross-spectrum between signals *x* and *y* over frequency *f*, with the expectation operator indicated as ⟨·⟩. The denominator denotes the square root of the product of the averaged power spectra of the signals *x* and *y* at the same frequency. Thus, COH between *x* and *y* is a cross-spectrum normalized by the square root of both signals’ power spectra product.

However, COH values can be contaminated by ‘Volume conduction’, a phenomenon in which surrounding potentials influence the measurement of the target’s electrical potential. Because EEG data often contain a mix of signals-of-interest and signals-of-no-interest, this can lead to spurious connectivity values when the two signals are not genuinely synchronized [19], [41], [42]. A significant challenge is that it is difficult to determine the extent to which the estimated value contains erroneous information. To address this issue, iCOH and PLV have been suggested.

**Imaginary Coherence** iCOH is obtained by projecting coherence onto an imaginary axis. iCOH discards certain estimated values by executing the projection assuming that volume conduction may affect the estimation [19], [43].

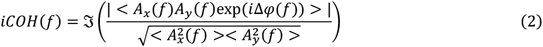

in which *ζ*(·) indicates the imaginary part.

When the data are contaminated by volume conduction, iCOH gives a more precise evaluation. However, it can discard valuable information about the synchronization between signals if volume conduction does not exist or is not that serious.

**Phase Locking Value** PLV normalizes amplitudes of Fourier-transformed signals to strictly phase synchronization between signals estimate [19], [44]. Thus, PLV is a version of COH with normalized amplitudes.

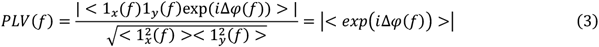

in which 1_*i*_ represents the normalized amplitude of signal *i*.

As each signal’s amplitudes have been normalized, the only remaining factor to consider is the phase difference. Theoretically, it may be reasonable, as a large amplitude product may overestimate the coherence value. However, the PLV estimate may not be correct because amplitudes reflect the signal-of-interest [19].

**Phase Slope Index** PSI measures unidirectional interactions between channels, and it estimates the consistency of phase slope across frequencies [38]. It calculates phase differences between adjacent frequencies based upon a predetermined bandwidth parameter. The phase difference is weighted by the two frequencies’ coherency. If the phase difference between frequencies is constantly changing and substantial coherence exists, the resulting value will deviate from 0. Thus, if the direction inferred from the phase difference remains constant across the given frequency range, it indicates the existence of distinct interactions between the signals. The sign of the phase difference reveals which signal is leading the other.

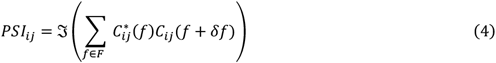

in which 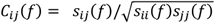 is the complex coherency, *s* is the cross-spectral matrix, *δf* is the frequency resolution, and *ζ*(·) denotes taking the imaginary part. *F* is the set of frequencies of interest.

PSI is known to be a robust measure against mixtures of independent sources. However, if the interaction between signals is not unidirectional, the estimation can fail to detect the interaction.

#### 2.3.2 The procedure to estimate Granger Causality in the time domain

We performed the same channel selection and data segmentation as in the frequency domain. However, we omitted the CAR step because of rank deficiency. Instead, we applied a moving average filter with a 5-point window to reduce noise after applying an interest band-pass filter. In addition, we downsampled the data from 512 Hz to 256 Hz to capture the data’s underlying trends better. We estimated the Bayesian information criterion to determine the best model order p. The trial matrices are concatenated to compute a coefficient matrix of the auto-regression (AR) model, which produces a single connectivity matrix for all trial matrices. To ensure that the AR model was stationary, we calculated the coefficient matrix’s spectral radius and confirmed that it was less than 1. For the statistical analysis, we performed Granger’s F-test, Block permutation test, and FDR with alpha = 0.05 [36]. The procedure is summarized in Fig. 2.

**Fig. 2.**
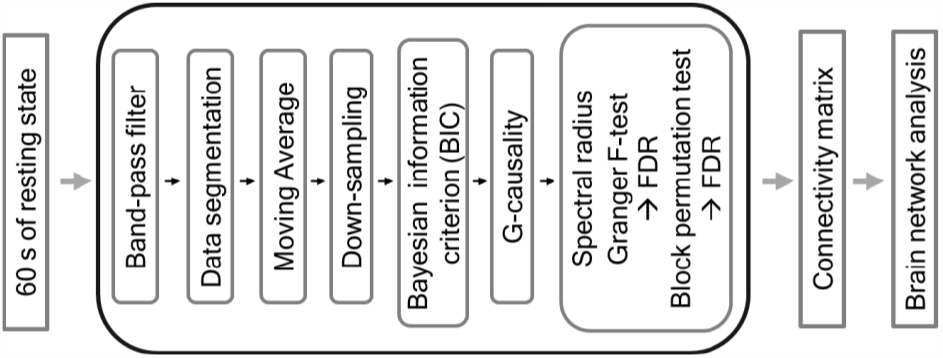
The procedure of estimation of Granger Causality in the time domain.

**Bivariate Granger Causality** To estimate causality between two channels, BVCG considers only the two channels for an auto-regression model [36], and it quantifies the causality by comparing the variance of each model’s prediction error. The random vector X_i_(*t*) contains time-series data of EEG channel i, in which X_i_(*t*) = [*x*_*i*_(*t*), *x*_*i*_(*t* − 1), …, *x*_*i*_(*t* − *p*)]. The random vector A_i_(*t*) is a coefficient vector of X_i_(*t*), in which A_i_(*t*) = [*a*_*i*_(*t*), *a*_*i*_(*t* − 1), …, *a*_*i*_(*t* − *p*)]. Finally, the random vector, E_i_, is a regression model’s error of the i^th^ channel, in which E_i_(*t*) = [*e*_*i*_(*t*), *e*_*i*_(*t* − 1), …, *e*_*i*_(*t* − *p*)].

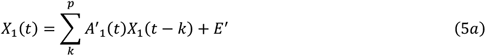

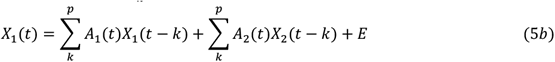

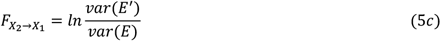

(5a) is a reduced model that omits a source variable X_2_, while (5b) is a full model that contains the source variable. Because of a regression model’s property, a reduced model will have a greater error as it contains fewer independent variables. Therefore, the natural logarithm of each error’s variance ratio must be greater than zero, known as G-causality [19], [35], [36]. (4c) is the G-causality of X_2_ to X_1_. If the full model’s error, E, is the same as the reduced model’s error, E′, G-causality is zero. Otherwise, G-causality is greater than zero, and a higher value indicates a greater causality.

**Multivariate Granger Causality** MVGC estimates the causality between two channels with all variables [38]. Theoretically, it can compute more precise G-causality because the regression model includes all possible source variables. However, it leads to overfitting and underestimates G-causality because unnecessary parameters may interrupt model fitting [19], [36]. Further, it has a very high computational cost.

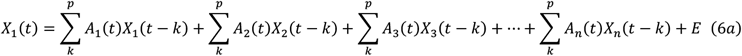

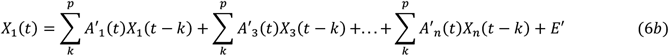

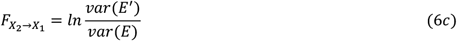

### 2.4 Brain Network Analysis

A network (or graph) is a set comprising nodes and edges. In the context of EEG data, the EEG channels can be considered nodes, and the connectivity between them can be considered edges. We measured graph features using the Brain Connectivity Toolbox [45]. We investigated resting state differences in the network’s efficiency and structure between BCI-illiteracy and -literacy. In this study, we limited ourselves to introducing the measures for analysis and its essential concept briefly. Readers interested in more details are referred to the papers referenced [22], [45].

#### 2.4.1 The measure of network efficiency

##### Strength

The strength of the connections between nodes in the network is represented by the weighted degrees. A network characterized by strong connections to each node suggests high network efficiency, as this typically correlates with more rapid information transfer between nodes compared to a network with weaker connections. The network’s average weighted degrees is a good measure to evaluate the network’s efficiency, which is called Strength (STR) [45], in which a higher value indicates greater efficiency.

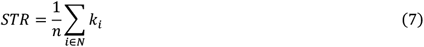

in which *k*_*i*_denotes a weighted degree of node *i*.

##### Clustering Coefficient

The network segregation reveals efficiency, as it represents the network’s ability to process specialized information [45]. Measuring how efficiently nodes in the network cluster with their neighbors based upon the number of triangles is an acceptable approach [45]–[48]. The Clustering Coefficient (CC) is a well-known measure of the network’s efficiency.

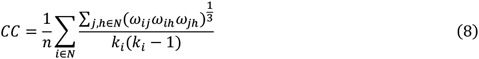

in which *w*_*ij*_ represents an edge between nodes *i*and *j*.

The denominator represents the number of triangles that involve node *i*, while the numerator corresponds to the number of all possible triples of node *i*. Then, the CC is measured as the average ratio of the number of all possible triples that a node can have to that of triangles that a node actually has in the network [45], [48]. It ranges between 0 and 1, in which higher values indicate greater efficiency.

##### Characteristic Path Length

The network’s integration can be measured by calculating the shortest path length between pairs of nodes in the network. When the shortest path length is small, it suggests that the information flow between pairs of nodes will be rapid and efficient. The Characteristic Path Length (CPL) quantifies network integration by calculating the average shortest path length between all pairs of nodes in the network [45]. CPL is not a ranged metric, and higher values indicate that the network is not integrated well.

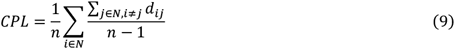

in which *d*_*ij*_ indicates the shortest path length between node *i*and *j*.

##### Global Efficiency

Global Efficiency (GE) measures the network’s integration by inversing the shortest path length [45], [47]. This makes sense because a shorter information flow path between nodes results in a better network. A GE value ranges between 0 and 1, and higher values indicate that the network is integrated well.

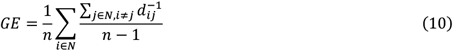

It is known generally that long paths affect CPL predominantly, while short paths affect GE predominantly by [45].

##### Local Efficiency

The network’s ‘fault tolerance’ is a notable example of efficiency. If the information flow involving node i remains intact even when this node is removed, owing to the compensatory role of its neighboring nodes, we can consider that the network is fault-tolerant[47]. To measure this property, it is necessary to examine the structure of the subgraphs within the network. Local Efficiency (LE) measures the average efficiency of the local subgraphs by estimating the sub-shortest path between nodes i, j, and h, which includes only the node i’s neighbors. The value ranges between 0 and 1, in which a higher value indicates better segregation.

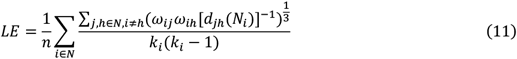

in which *d*_*jh*_(*N*_*i*_) represents the shortest path length that includes only the neighbors of node *i*.

#### 2.4.2 The measure of network structure

##### Betweenness Centrality

The hub node plays a crucial role in the network’s segregation and integration [45], [48], [49]. Identifying a hub node provides the key way to understand the network’s structure. Betweeness Centrality (BC) is one of the measures used to identify a hub node [50].

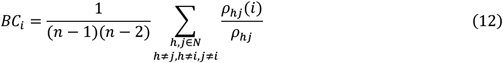

in which *p*_*hj*_ presents the number of shortest paths between nodes *h* and *j*, and *p*_*hj*_(*i*) indicates the number of shortest paths between h and j that pass through i.

BC quantifies the degree of a node’s hub status by measuring its involvement in the shortest paths [45], [48], [50] and is used to estimate modules in the network.

##### Modularity

Newman’s algorithm is a measure used widely to estimate the network’s modules. The essential mechanism is to remove the node with the highest BC value and calculate the ‘Modularity’. This process is repeated until the optimal point is reached and results in the identification of the best possible modules for the network [45], [51]–[53]. The modularity value falls within the range of -1 to 1, and most values fall between 0 and 1. A value near 0 suggests that the partition is occurring randomly, while a value nearer to 1 indicates a highly efficient and meaningful partition.

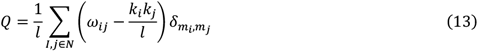

in which *l* indicates a sum of all weighted edges, *m*_*i*_ is the module that includes node *i*, and *δ*_*mi,mj*_ =1 if nodes *i*and *j* are in the same module, and 0 otherwise.

###### Z-score and Participation Coefficient

Once modules are determined, we can estimate the ‘Z-score’ and ‘Participation Coefficient (PC)’. The Z-score measures the degree to which a node is connected with nodes in another module [54]. A node with a higher value indicates that the node may possess a hub status as it is connected extensively within its module.

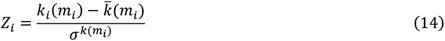

in which *k*(*m*_*i*_) represents the within-module weighted degree of node *i*, and 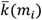 mean and σ^k(mi)^ standard deviation of the within-module m_i_ weighted degree distribution, respectively.

Unlike the Z-score, the PC measures the degree of a node’s hub status by measuring the extent to which a node is connected to different modules within the network [54].

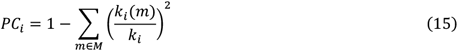

in which *M* is the set of modules in the network, and *k*_*i*_(*m*) indicates the weighted degree of node *i*that connects to nodes in module *m*.

## 3 Results

### 3.1 Results of BCI performance accuracy and categorization

The results of MI-BCI’s performance in each dataset are shown in Figure 3. In the primary dataset, there were 20 BCI-illiterates, while there were 16 BCI-illiterates. The average accuracy of BCI-literacy was 78.73% ± 7.03%, while that of BCI-illiteracy was 55.54% ± 3.05%. The mean of accuracy difference between 120 times validation for each subject in the BCI-literate group was 5.88% ± 1.50%, while it was 5.82% ± 0.52% in the BCI-illiterate group.

**Fig. 3.**
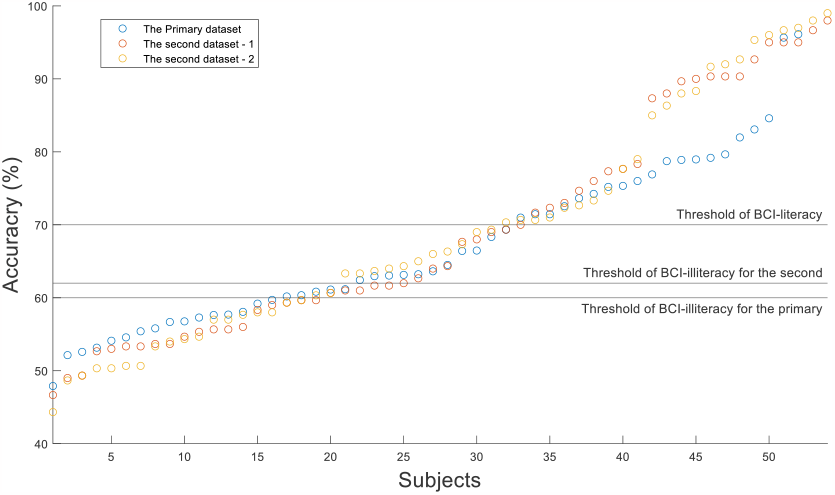
The performance results of MI-BCI from each dataset.

In the second dataset - 1, 22 subjects were categorized as BCI-literate, while 24 were categorized as BCI-illiterate. BCI-literate participants’ average accuracy was 84.97% ± 9.42%, and was 56.00% ± 4.25% for BCI-illiterate participants. The mean of accuracy difference between the validation for each subject was 5.97% ± 2.34% in the BCI-literate group and 8.32% ± 2.08% in the BCI-illiterate group.

In the second dataset - 2, 23 subjects were categorized as BCI-literate and 20 were categorized as BCI-illiterate. BCI-literate participants’ average accuracy was 84.27% ± 10.66%, while it was 54.42% ± 4.58% for BCI-illiterate participants. The mean of accuracy difference between the validation for each subject in the BCI-literate group was 6.38% ± 2.96% and 8.37% ± 1.63% in the BCI-illiterate group.

As indicated in Table 1, the accuracy difference between 120 times validation for each subject appears to be consistent between the two groups in the primary dataset. In contrast, this difference does not exhibit a similar pattern in the second dataset. Based upon this result, we infer that the grouping in the primary dataset was stable, while it was not in the secondary dataset.

**Table 1.**
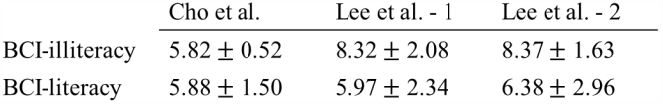
The mean of accuracy difference between 120 times validation for each subject.

### 3.2 The difference in network efficiency between BCI-literate and -illiterate groups in the primary dataset

Table 2 displays the mean and standard deviation of each graph feature for each metric across different frequency ranges for the BCI-literate and -illiterate groups (* indicates the values are scored by 10). The values that differed significantly between the two groups are highlighted in bold. Table 3 displays the p-values from the Wilcoxon rank-sum test and *t*-test, and highlights those below 0.05. In addition, Fig. 4. displays box plots with scatter plots for metrics in the frequency range that exhibits significant differences between the BCI-literate and -illiterate groups. As categorized by the whisker lengths, outliers were identified by a red cross, and no outliers were detected in COH, PLV, and MVGC. It was observed that regardless of the metric and frequency range, the BCI-literate group generally exhibited higher network efficiency than the BCI-illiterate group.

**Table 2.**
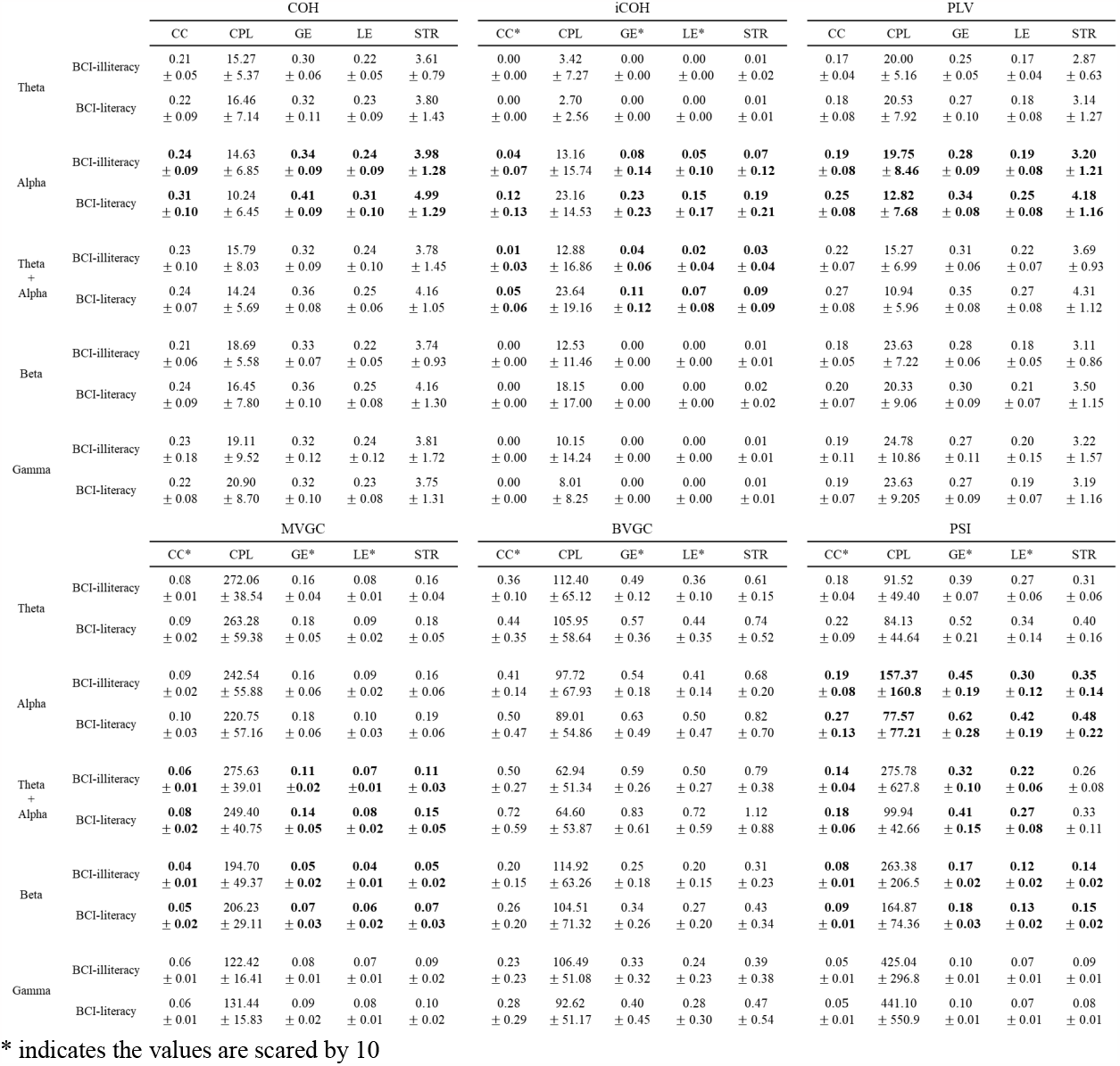
The statistic of each graph feature for each metric across different frequency ranges in the primary dataset.

**Table 3.**
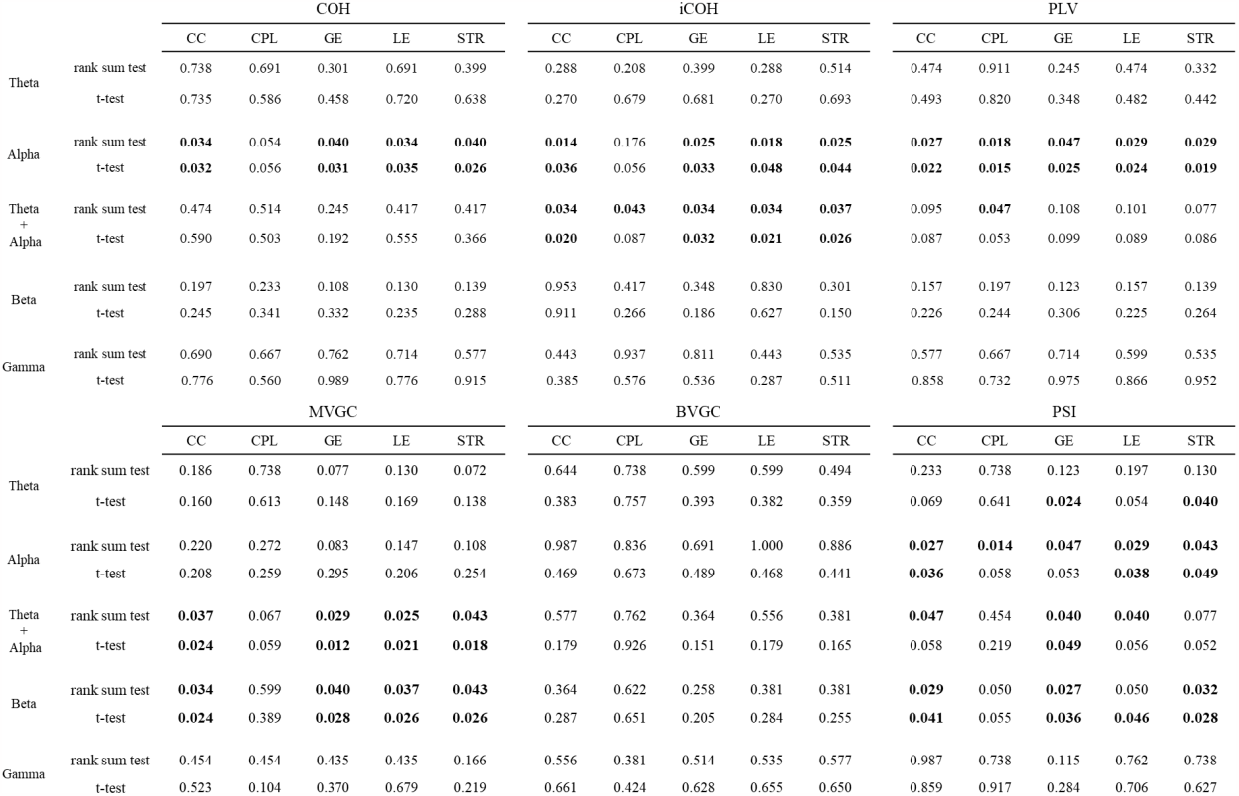
The p-values of each graph feature using all subjects in the primary dataset.

**Fig. 4.**
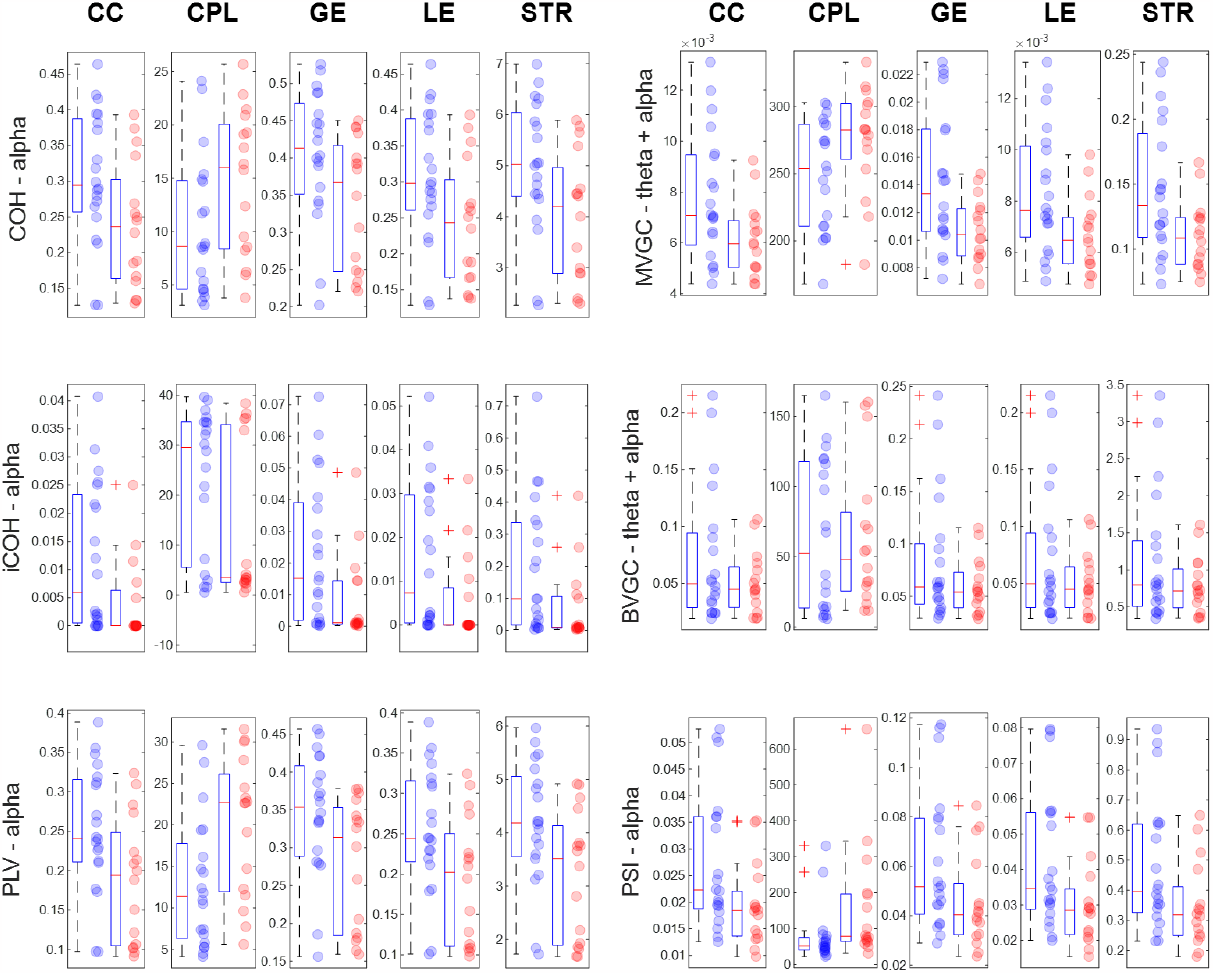
The box plots with scatter plots for metrics in the frequency range that exhibits significant differences between BCI-literate and illiterate groups from the primary dataset. The left bar and blue circle indicate BCI-literacy, while right bar and red circle refer to BCI-illiteracy.

However, with respect to the frequency range, FC and EC demonstrated different patterns. In the case of FC, the alpha band could be considered a crucial frequency range to evaluate connectivity in network analysis, regardless of the metric used, as summarized in Table 3. It is also interesting to observe that COH and PLV exhibited remarkably similar distributions across all graph features. However, only PLV showed a significant difference in CPL. As Table 2 shows, there was a significant difference in the theta + alpha exhibited by iCOH. However, it is worth noting that iCOH tended to yield much smaller values than COH and PLV because of the unique properties that we mentioned previously. Ultimately, this characteristic led to a wider distribution range. In the case of EC, the crucial frequency range for the evaluation differed depending upon the metric used. MVGC produced a significant difference in the theta + alpha and beta, while BVGC did not in all bands. PSI yielded a significant difference in alpha, theta + alpha, and beta. Further, only PSI showed a significant difference in CPL among EC.

### 3.3 The analysis of graph structure between BCI-literate and -illiterate groups

Fig. 5. displays the average connectivity matrices of each group’s subjects. It is evident that the connectivity matrix’s pattern remained similar for both the BCI-illiterate and literate groups within the same metrics. Although the iCOH matrix appeared to differ between the two groups, this is attributable to iCOH’s unique property, which causes a substantial deviation of the connectivity value within the groups.

**Fig. 5.**
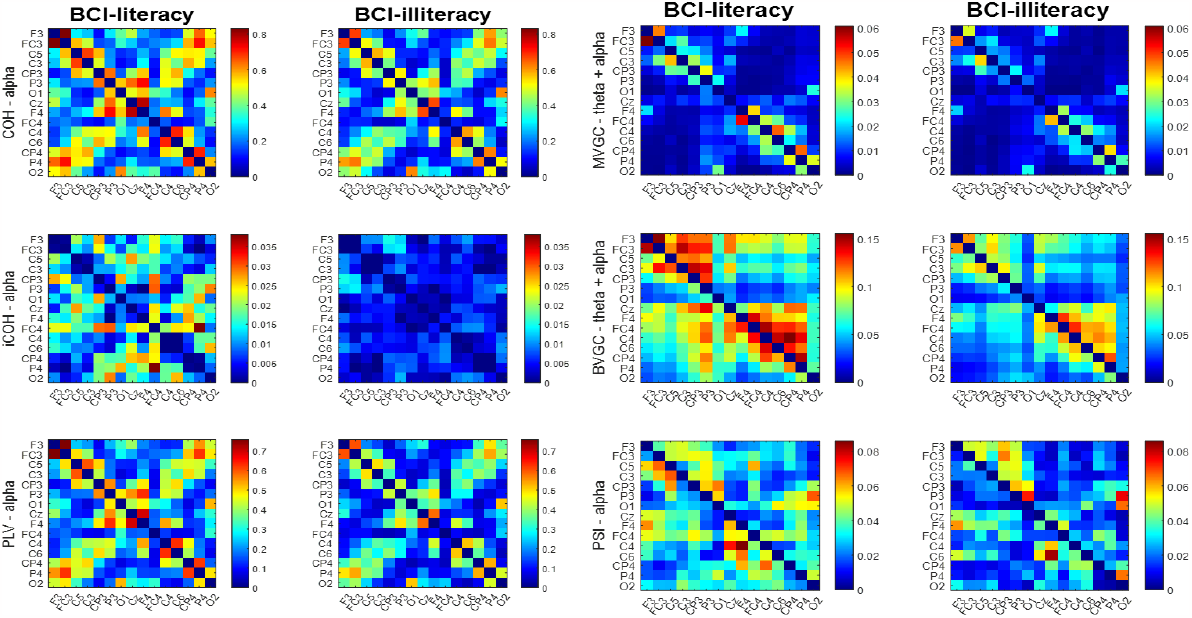
The average connectivity matrices of each group’s subjects in the primary dataset.

The major difference was observed in the connectivity value, in which the BCI-literate group displayed a stronger connectivity value. Table 4 shows to which modules the channels belong. However, the Q values suggested that the modular decomposition may result from chance rather than reflect noticeable graph features as they are too close to 0. In addition, in Fig. 6., it is observed that all channels are clustered together regardless of the metric and group, indicating that there is no hub node. Although PSI appeared to have a hub node, this was insignificant, as the Q value was zero. According to Fig 6., all metrics displayed a low BC, implying that no channel held a distinct node status.

**Table 4.**
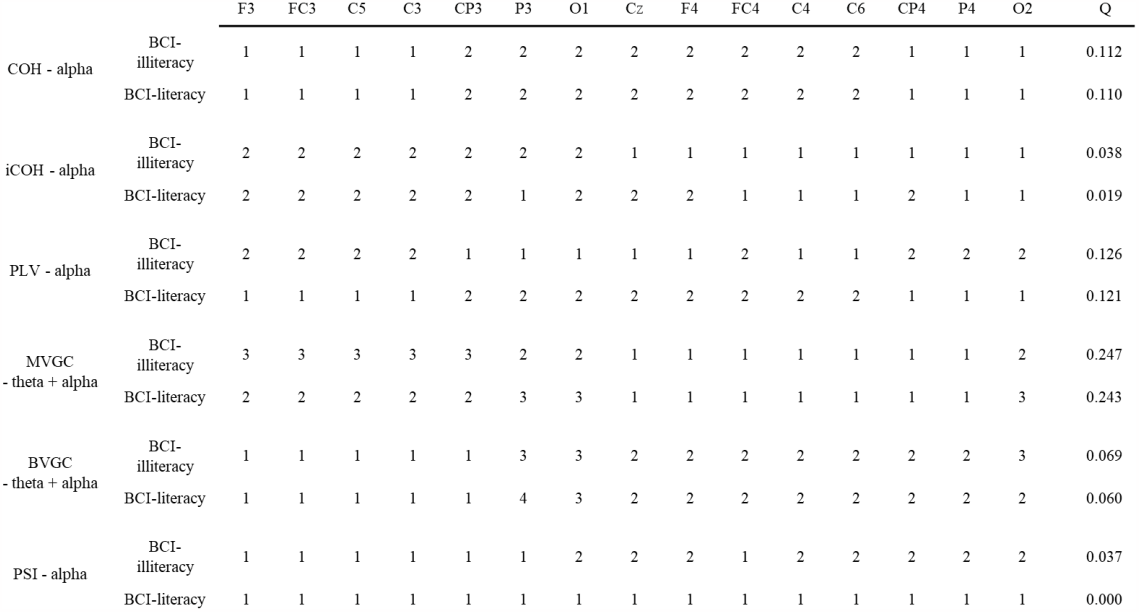
The modules that each channel belongs to and Q value.

|  |  | F3 | FC3 | C5 | C3 | CP3 | P3 | O1 | Cz | F4 | FC4 | C4 | C6 | CP4 | P4 | O2 | Q |
| --- | --- | --- | --- | --- | --- | --- | --- | --- | --- | --- | --- | --- | --- | --- | --- | --- | --- |
| COH - alpha | BCI-illiteracy | 1 | 1 | 1 | 1 | 2 | 2 | 2 | 2 | 2 | 2 | 2 | 2 | 1 | 1 | 1 | 0.112 |
|  | BCI-literacy | 1 | 1 | 1 | 1 | 2 | 2 | 2 | 2 | 2 | 2 | 2 | 2 | 1 | 1 | 1 | 0.110 |
| iCOH - alpha | BCI-illiteracy | 2 | 2 | 2 | 2 | 2 | 2 | 2 | 1 | 1 | 1 | 1 | 1 | 1 | 1 | 1 | 0.038 |
|  | BCI-literacy | 2 | 2 | 2 | 2 | 2 | 1 | 2 | 2 | 2 | 1 | 1 | 1 | 2 | 1 | 1 | 0.019 |
| PLV - alpha | BCI-illiteracy | 2 | 2 | 2 | 2 | 1 | 1 | 1 | 1 | 1 | 2 | 1 | 1 | 2 | 2 | 2 | 0.126 |
|  | BCI-literacy | 1 | 1 | 1 | 1 | 2 | 2 | 2 | 2 | 2 | 2 | 2 | 2 | 1 | 1 | 1 | 0.121 |
| MVGC - theta + alpha | BCI-illiteracy | 3 | 3 | 3 | 3 | 3 | 2 | 2 | 1 | 1 | 1 | 1 | 1 | 1 | 1 | 2 | 0.247 |
|  | BCI-literacy | 2 | 2 | 2 | 2 | 2 | 3 | 3 | 1 | 1 | 1 | 1 | 1 | 1 | 1 | 3 | 0.243 |
| BVGC - theta + alpha | BCI-illiteracy | 1 | 1 | 1 | 1 | 1 | 3 | 3 | 2 | 2 | 2 | 2 | 2 | 2 | 2 | 3 | 0.069 |
|  | BCI-literacy | 1 | 1 | 1 | 1 | 1 | 4 | 3 | 2 | 2 | 2 | 2 | 2 | 2 | 2 | 2 | 0.060 |
| PSI - alpha | BCI-illiteracy | 1 | 1 | 1 | 1 | 1 | 1 | 2 | 2 | 2 | 1 | 2 | 2 | 2 | 2 | 2 | 0.037 |
|  | BCI-literacy | 1 | 1 | 1 | 1 | 1 | 1 | 1 | 1 | 1 | 1 | 1 | 1 | 1 | 1 | 1 | 0.000 |

**Fig. 6.**
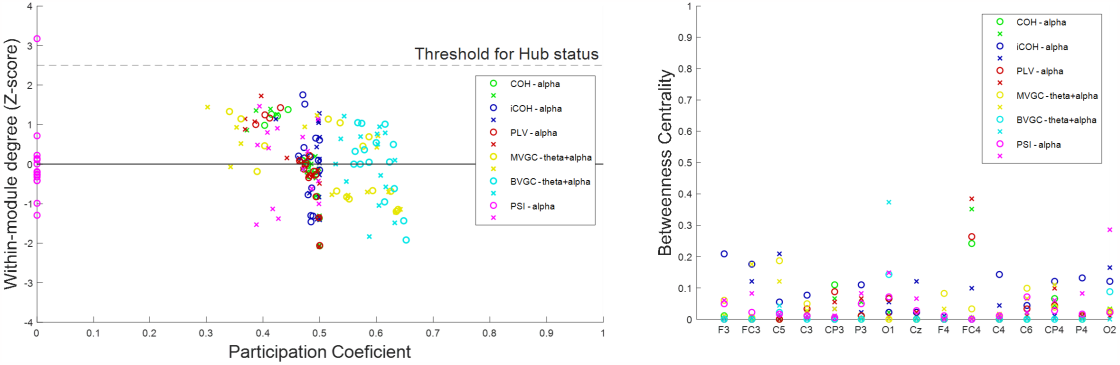
The distribution of PC, Z-score, and BC. Left figure indicates PC and Z-score of each metric within the most significant frequency range. Right figure exhibits BC values for each channel, as derived from each metric.

### 3.4 The correlation between network efficiency and BCI performance

We investigated the correlation between network efficiency and BCI performance in all subjects, focusing on the metrics and frequency bands that displayed significant differences between the two groups. Fig. 7. displays the least-squares lines derived from the results, without outliers. Table 5. shows the results’ correlation value, r, and corresponding p-value, in which the highest correlation is highlighted in bold. While COH and PLV exhibited similar results, PLV yielded slightly higher values. Further, in the case of PLV, the highest correlation was observed with STR, while it was with CC for COH. Except for CPL, EC showed a higher correlation than FC. In particular, MVGC had the highest correlation (r=0.38) with GE among the 4 metrics. However, if the outliers were excluded, the CC of PSI had the highest correlation (r=0.40).

**Table 5.** The correlation between graph features and BCI performance of the metrics in the frequency range that exhibited the significant difference in the primary dataset.

|  |  | CC | CPL | GE | LE | STR |
| --- | --- | --- | --- | --- | --- | --- |
| COH - alpha | r-value | **0.290** | -0.276 | 0.271 | 0.285 | 0.289 |
|  | p-value | **0.037** | 0.048 | 0.052 | 0.041 | 0.038 |
| PLV - alpha | r-value | 0.306 | -0.302 | 0.292 | 0.303 | **0.307** |
|  | p-value | 0.027 | 0.030 | 0.035 | 0.029 | **0.027** |
| MVGC<br>- theta + alpha | r-value | 0.312 | -0.206 | **0.383** | 0.316 | 0.340 |
|  | p-value | 0.024 | 0.143 | **0.005** | 0.022 | 0.014 |
| PSI - alpha | r-value | **0.342** | -0.268 | 0.323 | 0.339 | 0.327 |
|  | p-value | **0.013** | 0.055 | 0.020 | 0.014 | 0.018 |

**Fig. 7.**
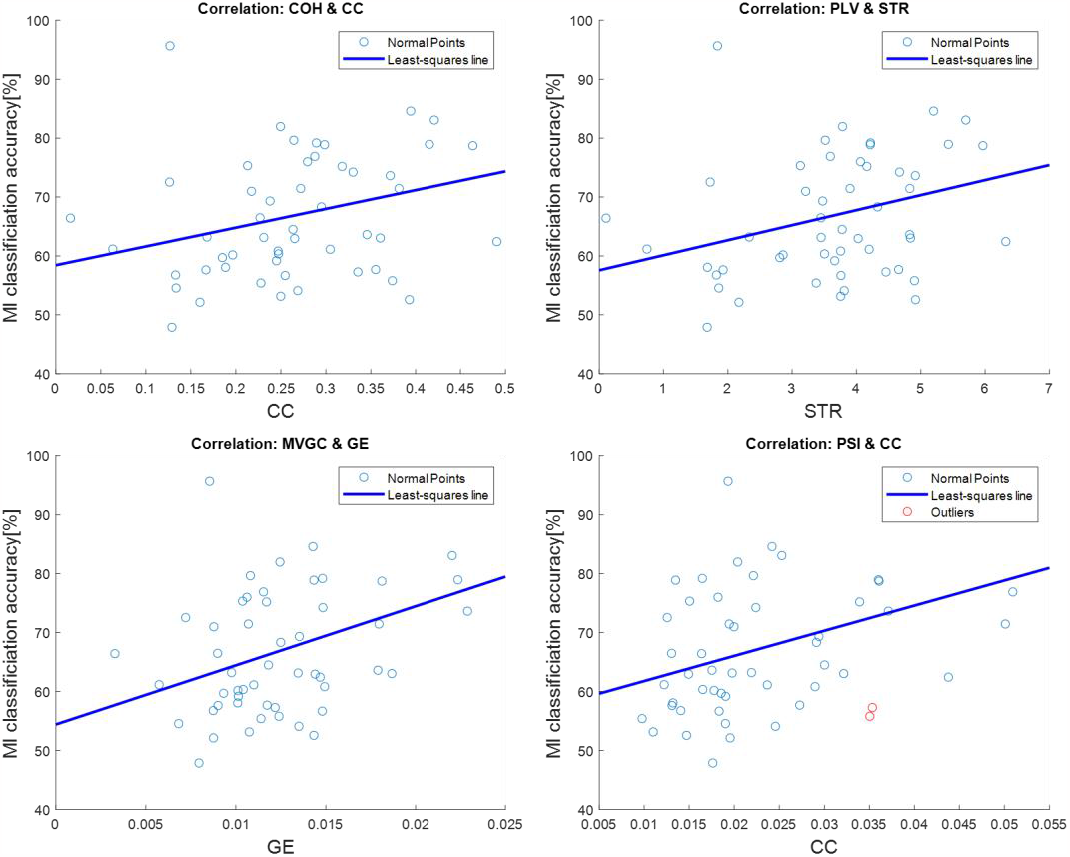
The least-square line of correlation between the graph features and MI-performance of each metric that showed the most significant difference in the primary dataset.

**Fig. 8.**
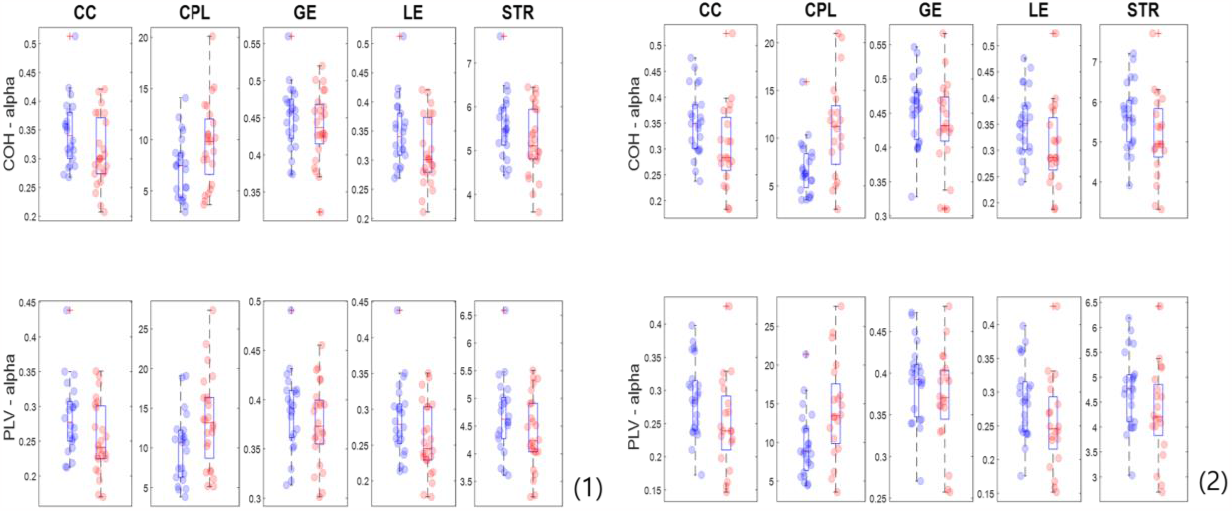
The box plot with scatter plot of COH and PLV from the second dataset. (1) corresponds to results derived from the second dataset -1, while (2) refers to the second dataset - 2. The left bar and blue circle mean BCI-literacy, while right bar and red circle denote BCI-illiteracy.

### 3.5 Replication of the results using the second dataset

We replicated the investigation of the primary dataset using the second dataset, which comprises two sections. The statistical results of the replication are presented in Table 6. (** indicates the values are scared by 10^2^.) In both sections, only FC exhibited a significant difference in the graph features between the two groups. Interestingly, COH and PLV differed significantly in CPL in both sections, unlike the result in the primary dataset. In particular, CC of COH produced significant differences in section 2. While not all graph features showed significant differences between the two groups, we did observe that the BCI-literate group exhibited consistently better network efficiency. Table 7 displays the correlation between BCI performance and graph features. The r value is in bold when p < 0.05. The correlation between BCI performance and COH’s CPL showed the highest correlation (r = -0.42). The least-squares lines without outliers are depicted in Fig. 9. The correlation values for COH and PLV were r = -0.44 and r = -0.37, respectively. Fig. 10 displays the average connectivity matrices of each group’s subjects in the second dataset, using COH, PLV, MVGC, and PSI. The matrices showed a pattern similar to the results of the primary dataset for COH, PLV, and MVGC, but differed for PSI.

**Table 6.** The statistical outcomes of the replication with the second dataset - 1,2. (1) details the statistics for each graph feature. (2) displays the p-values for each graph feature using all subjects.

| (1) |  | The second dataset - 1 |  |  |  |  | The second dataset - 2 |  |  |  |  |
| --- | --- | --- | --- | --- | --- | --- | --- | --- | --- | --- | --- |
|  |  | CC | CPL | GE | LE | STR | CC | CPL | GE | LE | STR |
| COH - alpha | BCI-illiteracy | 0.31<br>± 0.06 | **9.61**<br>± **4.03** | 0.44<br>± 0.05 | 0.31<br>± 0.06 | 5.21<br>± 0.79 | **0.30**<br>± **0.08** | **11.07**<br>± **5.12** | 0.43<br>± 0.06 | **0.31**<br>± **0.08** | 5.13<br>± 1.05 |
|  | BCI-literacy | 0.34<br>± 0.06 | **7.30**<br>± **3.05** | 0.45<br>± 0.04 | 0.35<br>± 0.06 | 5.56<br>± 0.74 | **0.31**<br>± **0.06** | **6.79**<br>± **2.82** | 0.45<br>± 0.05 | **0.35**<br>± **0.06** | 5.65<br>± 0.82 |
| PLV - alpha | BCI-illiteracy | 0.25<br>± 0.05 | **13.40**<br>± **5.77** | 0.38<br>± 0.04 | 0.26<br>± 0.05 | 4.37<br>± 0.67 | 0.25<br>± 0.07 | **14.24**<br>± **6.36** | 0.37<br>± 0.06 | 0.25<br>± 0.07 | 4.29<br>± 0.92 |
|  | BCI-literacy | 0.28<br>± 0.05 | **10.02**<br>± **4.34** | 0.39<br>± 0.04 | 0.29<br>± 0.05 | 4.68<br>± 0.69 | 0.29<br>± 0.06 | **9.57**<br>± **4.23** | 0.39<br>± 0.05 | 0.29<br>± 0.06 | 4.74<br>± 0.75 |
| MVGC**<br>- theta + alpha | BCI-illiteracy | 0.57<br>± 0.41 | 214.7<br>± 57.2 | 0.79<br>± 0.51 | 0.64<br>± 0.04 | 8.86<br>± 6.73 | 0.52<br>± 0.26 | 229.4<br>± 49.1 | 0.75<br>± 0.40 | 0.59<br>± 0.27 | 8.44<br>± 4.68 |
|  | BCI-literacy | 0.56<br>± 0.20 | 218.9<br>± 72.6 | 0.75<br>± 0.26 | 0.61<br>± 0.20 | 8.51<br>± 3.17 | 0.52<br>± 0.15 | 238.6<br>± 40.8 | 0.78<br>± 0.24 | 0.58<br>± 0.14 | 8.66<br>± 2.66 |
| PSI* - alpha | BCI-illiteracy | 0.24<br>± 0.10 | 97.97<br>± 152 | 0.54<br>± 0.23 | 0.37<br>± 0.16 | 0.43<br>± 0.17 | 0.30<br>± 0.27 | 72.04<br>± 36.1 | 0.66<br>± 0.53 | 0.47<br>± 0.41 | 0.53<br>± 0.43 |
|  | BCI-literacy | 0.28<br>± 0.12 | 16.45<br>± 38.2 | 0.63<br>± 0.28 | 0.43<br>± 0.19 | 0.50<br>± 0.21 | 0.30<br>± 0.12 | 59.52<br>± 31.6 | 0.68<br>± 0.25 | 0.46<br>± 0.18 | 0.53<br>± 0.20 |
| (2) |  | The second dataset - 1 |  |  |  |  | The second dataset - 2 |  |  |  |  |
|  |  | CC | CPL | GE | LE | STR | CC | CPL | GE | LE | STR |
| COH - alpha | rank sum test | 0.066 | **0.038** | 0.448 | 0.073 | 0.163 | **0.031** | **0.003** | 0.336 | **0.031** | 0.066 |
|  | t-test | 0.058 | **0.034** | 0.353 | 0.063 | 0.123 | **0.039** | **0.001** | 0.282 | **0.047** | 0.079 |
| PLV - alpha | rank sum test | 0.070 | **0.030** | 0.422 | 0.077 | 0.169 | 0.053 | **0.008** | 0.336 | 0.059 | 0.111 |
|  | t-test | 0.058 | **0.031** | 0.375 | 0.065 | 0.133 | 0.052 | **0.007** | 0.234 | 0.062 | 0.086 |
| MVGC<br>- theta + alpha | rank sum test | 0.461 | 0.590 | 0.590 | 0.700 | 0.410 | 0.247 | 0.670 | 0.258 | 0.201 | 0.219 |
|  | t-test | 0.858 | 0.829 | 0.758 | 0.757 | 0.824 | 0.988 | 0.505 | 0.777 | 0.938 | 0.852 |
| PSI - alpha | rank sum test | 0.248 | 0.502 | 0.222 | 0.222 | 0.231 | 0.268 | 0.184 | 0.128 | 0.228 | 0.210 |
|  | t-test | 0.316 | 0.311 | 0.223 | 0.280 | 0.254 | 0.894 | 0.232 | 0.919 | 0.914 | 0.986 |

\*\* indicates the values are scared by $10^2$ .

**Table 7.** The correlation between graph features and BCI performance in the second dataset.

|  |  | The second dataset - 1 |  |  |  |  | The second dataset - 2 |  |  |  |  |
| --- | --- | --- | --- | --- | --- | --- | --- | --- | --- | --- | --- |
|  |  | CC | CPL | GE | LE | STR | CC | CPL | GE | LE | STR |
| COH - alpha | r-value | 0.223 | -0.263 | 0.218 | 0.102 | 0.182 | 0.278 | **-0.421** | 0.268 | 0.176 | 0.248 |
|  | p-value | 0.106 | 0.055 | 0.113 | 0.464 | 0.188 | 0.042 | **0.002** | 0.050 | 0.203 | 0.071 |
| PLV - alpha | r-value | 0.215 | -0.238 | 0.209 | 0.097 | 0.172 | 0.257 | **-0.354** | 0.247 | 0.182 | 0.238 |
|  | p-value | 0.118 | 0.083 | 0.129 | 0.486 | 0.212 | 0.061 | **0.009** | 0.072 | 0.189 | 0.084 |

**Fig. 9.**
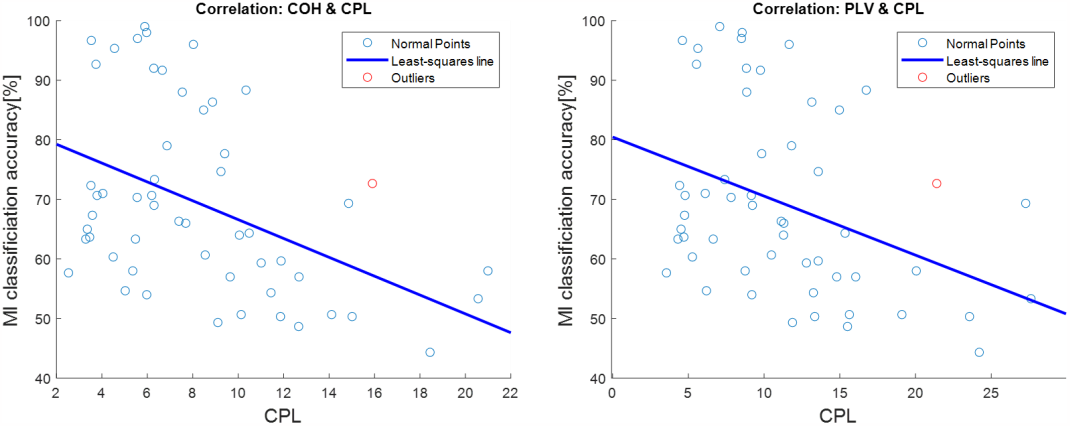
The least-square line of correlation between the graph features and BCI performance of COH and PLV in the second dataset – 2.

**Fig. 10.**
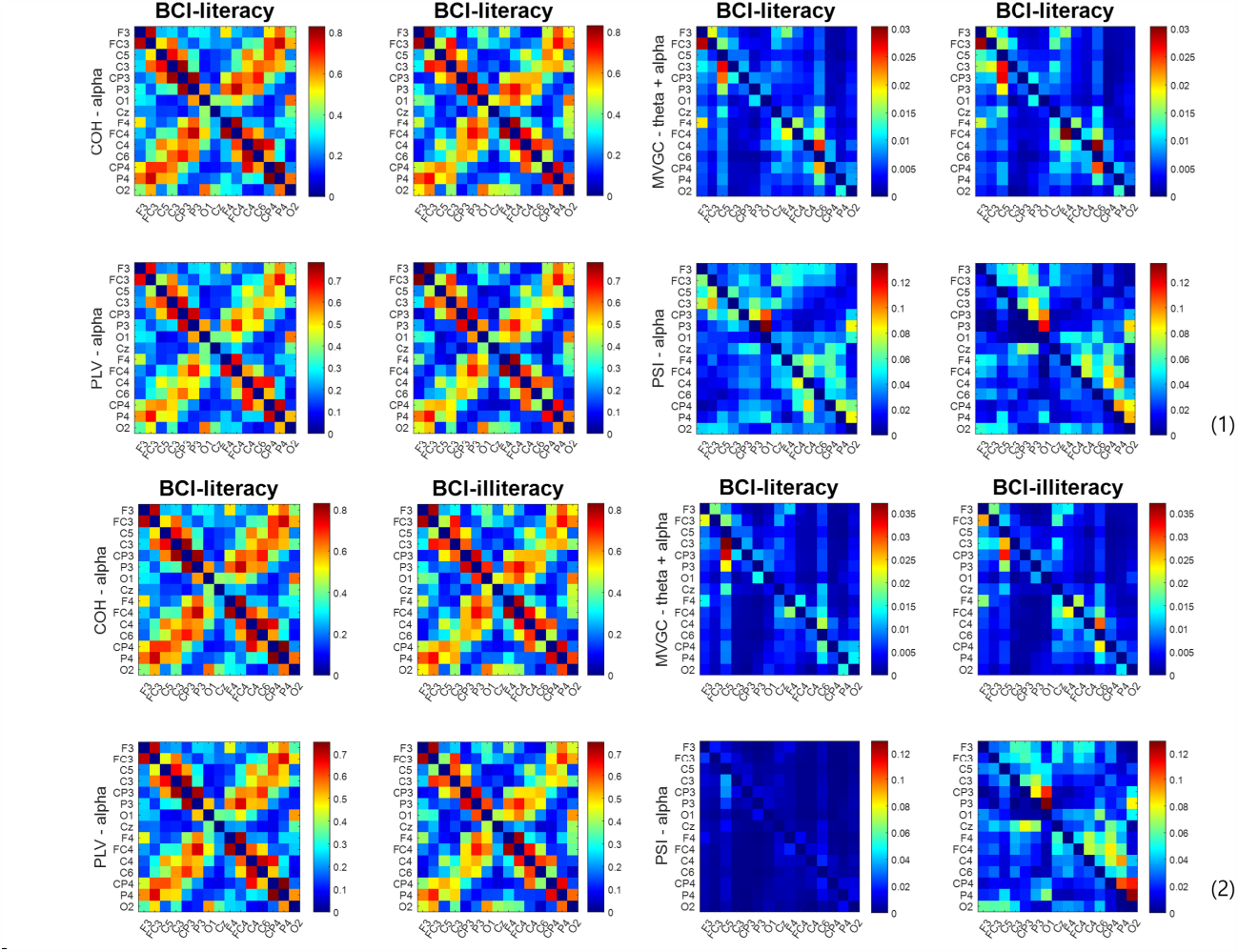
The average connectivity matrices of each group’s subjects in the second dataset – 1,2. (1) indicates the results from the second dataset -1, whereas (2) is from the second dataset - 2.

### 3.6 The results of the integrated dataset

Each group’s subjects’ average connectivity matrices in the integrated dataset using COH and PLV are presented in Fig. 11. These results resembled those obtained from the individual datasets closely. Table 8 provides a summary of the statistical outcomes from the integrated dataset. All graph features showed significant differences between the two groups. Among the five features, the CPL stood out as the most distinctive, with p-value = 0. In addition, we conducted a bootstrap test between the BCI-illiterate and -literate groups. To do so, we selected 25 subjects from each group randomly and performed 1000 iterations. All features’ p-values were 0. We performed the Kolmogorov-Smirnov test to check the features’ normality, which indicated that all features in both groups were distributed normally. We present the least square line of correlation between BCI performance and CPL in Fig. 12. The r values for COH and PLV were -0.31 and -0.30 with p-value = 0, respectively. Fig. 13. illustrates the distribution of graph features from the integrated dataset, as well as those from the primary and second dataset - 1,2. The results showed that the BCI-literacy’s network efficiency was significantly higher than that of BCI-illiteracy. The primary dataset had a wider gap between BCI-illiteracy and -literacy, while the difference was more subtle in the other datasets. However, subjects within each group were clustered together more closely in the second dataset.

**Fig. 11.**
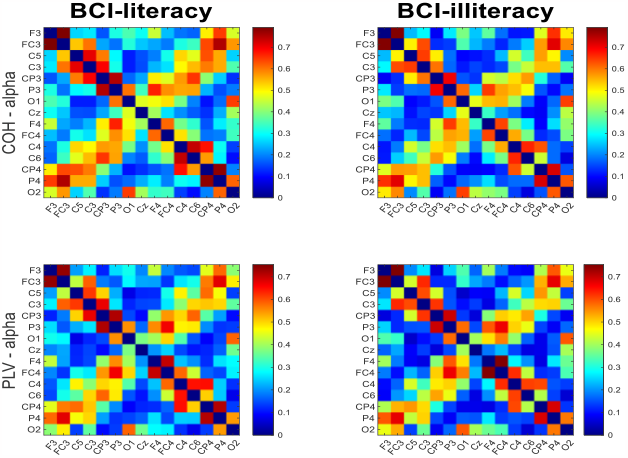
The average connectivity matrices of each group’s subjects in the integrated dataset.

**Fig. 12.**
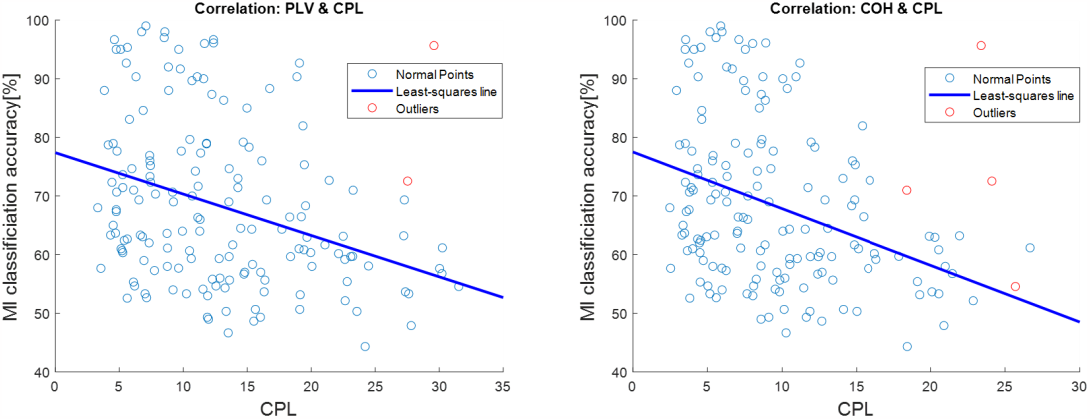
The least square line of correlation between CPL and BCI performance of COH and PLV in the integrated dataset.

**Fig. 13.**
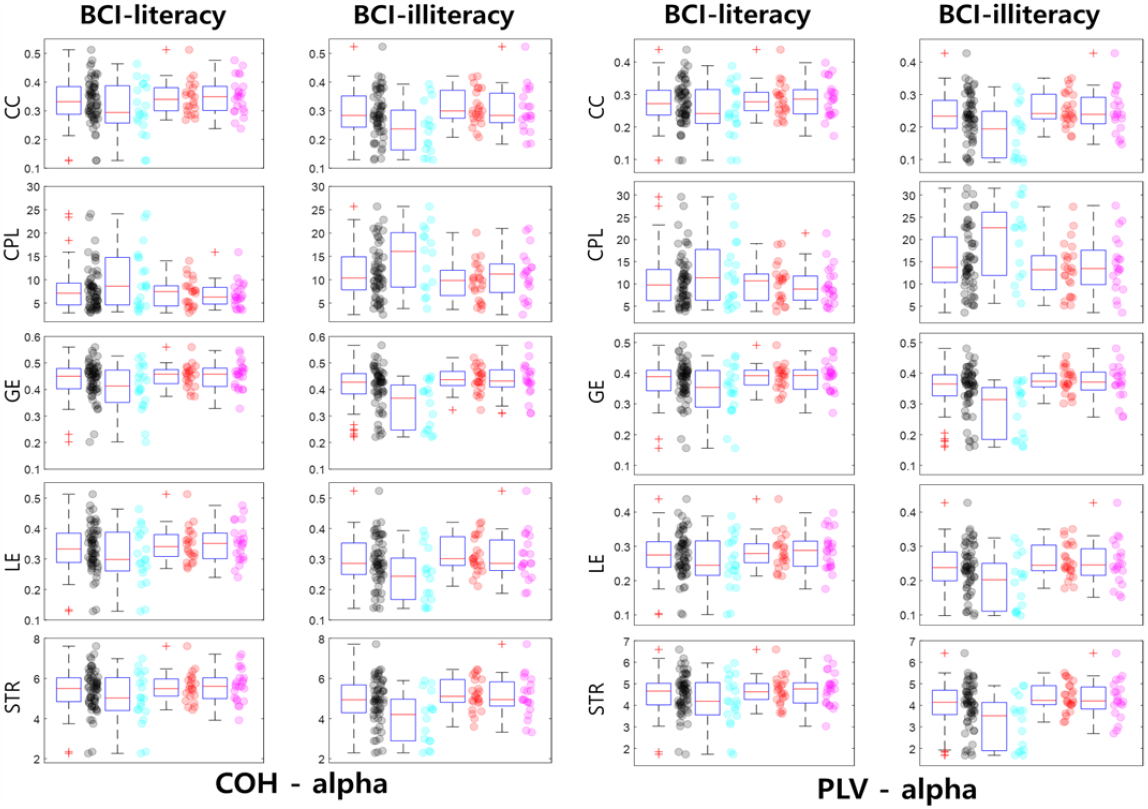
The box plots with scatter plots for COH and PLV in alpha range from the all dataset. The first bar and black circle, the second bar and cyan circle, the third bar and red circle, and the last bar and pink circle indicate the integrated, primary, second – 1, and second -2 dataset, respectively.

## 4 Discussion

### 4.1 The variation in estimating connectivity across different metrics and datasets

Each metric measures connectivity according to its unique perspective and hypothesis [19]. As a result, the estimated connectivity varies based upon the metric used. However, interestingly, the difference in network efficiency values between BCI-literacy and -illiteracy, estimated by diverse graph theory measures, was similar regardless of the metric used. The BCI-literacy’s resting-state EEG network demonstrated higher network efficiency compared to BCI-illiteracy. Thus, although the network’s pattern and connections changed according to metrics, its efficiency remained constant.

In the primary dataset, most metrics exhibited a significant difference between the two groups, while BVGC did not. This is because other sources contaminated BVGC’s G-causality [19], [35], [36]. While iCOH showed a significant difference, most of the estimated graph feature’s values were zero as iCOH discards too many connectivity values when the volume conduction is weak [19], [43]. Although PLV was found to have a higher correlation value than COH in the primary dataset, the converse was observed in other datasets. This may be attributable to the variation in the amplitude and degree of volume conduction in the dataset. These results are consistent with what we anticipated theoretically in the previous section. Hence, these factors should be considered during the analysis.

In the primary dataset, EC estimated resting-state connectivity better than FC when identifying BCI-illiteracy. PSI estimated connectivity well without a permutation test, demonstrating robustness to spurious connectivity as Nolte et al. claimed [38]. In addition, it is worth noting that capturing unidirectional or bidirectional information flow resulted in good estimations of both. However, this result was valid only in the primary dataset.

In the second dataset and the integrated dataset, only COH and PLV in the alpha band were appropriate metrics. The results of EC may be relevant only to the primary dataset. However, given that the categorization did not perform well in the second dataset because of the insufficient number of trials, the primary dataset’s result would be more accurate. Most importantly, as the two datasets’ settings and goals differed, a fair comparison of the results was not possible.

Nevertheless, we observed that BCI-literacy tends to have better network efficiency during the resting-state, despite the differences in the metrics and datasets. This tendency has been observed commonly in previous studies as well [23], [25]. We concluded that each metric’s hypothesis and perspective are appropriate to capture the information flow in the brain in specific situations and with specific assumptions. Therefore, it seems valid to predict BCI performance with resting-state connectivity.

### 4.2 Inappropriate interpretation of connectivity

From our findings, we claimed that predicting BCI performance based upon the connection between specific brain regions may be inappropriate. Lee et al. proposed that the strength of the connection between the supplementary motor area (SMA) and dorsolateral prefrontal cortex (DLPFC) during the resting-state can be indicative of BCI performance [25]. This is because several studies have shown that these two regions are involved actively during motor-related or cognitive tasks [55]–[60], and the authors found a stronger connection between the two regions in the BCI-literate group during the resting-state. However, they used only ‘Dynamic Causal Modeling’ to estimate connectivity, which implies that their results may vary dramatically when other metrics are employed. Moreover, it is not logically sound to conclude that these two regions’ higher activation during the motor-related task results from their stronger connection in the resting state. Rather, it could be the result of the strong connection between two very different regions during the resting state, considering numerous studies on DMN [27]–[29]. Thus, researchers need to be cautious not to draw incorrect conclusions from a connectivity result.

### 4.3 The proper frequency ranges for connectivity metrics to identify BCI-illiteracy

To identify BCI-illiteracy, the appropriate frequency range varies depending upon the metrics. For FC, the alpha band is the appropriate frequency range. However, in the case of EC, alpha, theta + alpha, and beta were found to be appropriate for the analysis. It can be inferred that estimating connectivity based upon phase differences between channels should be restricted to the alpha range. Although Zhan et al. reported that COH’s resting-state connectivity in 4-14 Hz exhibited a significant difference between BCI-literate and illiterate groups, the difference may be more pronounced if the analysis is restricted to the alpha band. While PSI is a metric in the frequency domain as well, it produces different results. It may indicate that calculating the phase difference between neighbouring frequency bins captures the information flow between channels more accurately or needs not to be restricted to the alpha range. When analyzing GC in the time domain, relying solely on the alpha range may not provide sufficient information to estimate the directional interaction between channels accurately. On the other hand, including too broad a range of frequencies (such as theta, alpha, and beta: 3-30 Hz) led to improper estimation as well.

Our interpretation was based solely upon the observation of the results. However, at present, it is still unclear why each metric requires a different frequency range to identify BCI-illiteracy with the resting-state. Therefore, further research is necessary to explore the relation between connectivity metrics and the frequency range to address our inference definitively.

### 4.4 The structure of the resting-state EEG connectivity

We investigated the network structure of resting-state EEG. In particular, we probed a hub node during the resting-state. It was found that neither the BCI-literate nor -illiterate groups had a hub node in all datasets. Interestingly, the structure was very similar between the two groups. The only difference was the strength of the connection between channels. In particular, COH and PLV’s connectivity matrices were remarkably similar across different datasets. These findings may suggest that coherence (COH) and phase locking value (PLV) are reliable metrics with which to assess connectivity. Alternatively, given that the default mode network (DMN) tends to be similar across individuals, these results may be somewhat expected.

### 4.5 Limitations

Estimating EEG connectivity is susceptible to the choice of EEG channels, indicating that our results may vary despite the application of rigorous statistical tests. This suggests that there may be an optimal channel to select for resting-state EEG connectivity, which makes channel selection a critical factor in the analysis. However, in this study, we did not investigate the optimal channel configuration, and adhered instead to the protocol established in a prior study [23]. Future research is warranted to identify the most effective channels for a resting-state EEG connectivity analysis.

It is currently uncertain whether directional information is helpful to estimate connectivity. Although it appeared to be useful in the primary dataset, it did not prove to be so in other datasets. Despite the fact that the second dataset did not have sufficient trials to classify BCI-literacy and -illiteracy accurately, the pivotal cause of this remains unknown. Therefore, further research is required to evaluate the directional information’s worth.

## 5. Conclusion

In this study, we examined the way that connectivity differed based upon metrics, frequency ranges, and datasets by analyzing resting-state EEG connectivity to provide a comprehensive understanding and insights into connectivity. In addition, we investigated which connectivity metric is the most reliable for identifying BCI-illiteracy by inspecting the correlation between BCI performance and graph features. For these, we assumed that BCI-literacy and -illiteracy’s resting-state EEG differs based upon previous studies. We used three different MI-BCI datasets and tested three FC and three EC metrics. Our observations showed that the actual results were some extent consistent with the theoretical expectation, and each metric had appropriate frequency ranges to identify BCI-illiteracy. While EC exhibited the highest correlation in the primary dataset, FC showed a robust estimation across the entire dataset. This suggested that the dataset affected the estimation of the metric’s connectivity. Based upon these results, we concluded that each metric has an appropriate hypothesis and perspective to measure connectivity in a specific context.

## Acknowledgments

This work was supported by the IITP (Institute of Information and Communications Technology Planning & Evaluation) grants funded by the Korea government (No. 2017-0-00451; No. 2019-0-01842).

## References

[1] J. R. Wolpaw, N. Birbaumer, D. J. McFarland, G. Pfurtscheller, and T. M. Vaughan, “Brain–computer interfaces for communication and control,” Clinical Neurophysiology, vol. 113, no. 6, pp. 767–791, Jun. 2002, doi: 10.1016/S1388-2457(02)00057-3.

[2] B. J. Edelman et al., “Noninvasive neuroimaging enhances continuous neural tracking for robotic device control,” Sci. Robot., vol. 4, no. 31, p. eaaw6844, Jun. 2019, doi: 10.1126/scirobotics.aaw6844.

[3] B. Blankertz, R. Tomioka, S. Lemm, M. Kawanabe, and K. Muller, “Optimizing Spatial filters for Robust EEG Single-Trial Analysis,” IEEE Signal Process. Mag., vol. 25, no. 1, pp. 41–56, 2008, doi: 10.1109/MSP.2008.4408441.

[4] A. Kübler, N. Neumann, J. Kaiser, B. Kotchoubey, T. Hinterberger, and N. P. Birbaumer, “Brain-computer communication: Self-regulation of slow cortical potentials for verbal communication,” Archives of Physical Medicine and Rehabilitation, vol. 82, no. 11, pp. 1533–1539, Nov. 2001, doi: 10.1053/apmr.2001.26621.

[5] J. Perelmouter and N. Birbaumer, “A binary spelling interface with random errors,” IEEE Trans. Rehab. Eng., vol. 8, no. 2, pp. 227–232, Jun. 2000, doi: 10.1109/86.847824.

[6] N. Birbaumer et al., “A spelling device for the paralysed,” Nature, vol. 398, no. 6725, pp. 297–298, Mar. 1999, doi: 10.1038/18581.

[7] A. Kübler and N. Birbaumer, “Brain–computer interfaces and communication in paralysis: Extinction of goal directed thinking in completely paralysed patients?,” Clinical Neurophysiology, vol. 119, no. 11, pp. 2658–2666, Nov. 2008, doi: 10.1016/j.clinph.2008.06.019.

[8] A. Kübler, N. Neumann, B. Wilhelm, T. Hinterberger, and N. Birbaumer, “Predictability of Brain-Computer Communication,” Journal of Psychophysiology, vol. 18, no. 2/3, pp. 121–129, Jan. 2004, doi: 10.1027/0269-8803.18.23.121.

[9] B. Z. Allison and C. Neuper, “Could Anyone Use a BCI?,” in Brain-Computer Interfaces, D. S. Tan and A. Nijholt, Eds., in Human-Computer Interaction Series., London: Springer London, 2010, pp. 35–54. doi: 10.1007/978-1-84996-272-8_3.

[10] C. Vidaurre and B. Blankertz, “Towards a Cure for BCI Illiteracy,” Brain Topogr, vol. 23, no. 2, pp. 194–198, Jun. 2010, doi: 10.1007/s10548-009-0121-6.

[11] M. C. Thompson, “Critiquing the Concept of BCI Illiteracy,” Sci Eng Ethics, vol. 25, no. 4, pp. 1217–1233, Aug. 2019, doi: 10.1007/s11948-018-0061-1.

[12] B. Blankertz et al., “Neurophysiological predictor of SMR-based BCI performance,” NeuroImage, vol. 51, no. 4, pp. 1303–1309, Jul. 2010, doi: 10.1016/j.neuroimage.2010.03.022.

[13] G. Pfurtscheller and F. H. Lopes Da Silva, “Event-related EEG/MEG synchronization and desynchronization: basic principles,” Clinical Neurophysiology, vol. 110, no. 11, pp. 1842–1857, Nov. 1999, doi: 10.1016/S1388-2457(99)00141-8.

[14] M. Ahn, H. Cho, S. Ahn, and S. C. Jun, “High Theta and Low Alpha Powers May Be Indicative of BCI-Illiteracy in Motor Imagery,” PLoS ONE, vol. 8, no. 11, p. e80886, Nov. 2013, doi: 10.1371/journal.pone.0080886.

[15] M. Kwon, H. Cho, K. Won, M. Ahn, and S. C. Jun, “Use of Both Eyes-Open and Eyes-Closed Resting States May Yield a More Robust Predictor of Motor Imagery BCI Performance,” Electronics, vol. 9, no. 4, p. 690, Apr. 2020, doi: 10.3390/electronics9040690.

[16] M. Grosse-Wentrup, B. Schölkopf, and J. Hill, “Causal influence of gamma oscillations on the sensorimotor rhythm,” NeuroImage, vol. 56, no. 2, pp. 837–842, May 2011, doi: 10.1016/j.neuroimage.2010.04.265.

[17] R. Zhang et al., “Predicting Inter-session Performance of SMR-Based Brain–Computer Interface Using the Spectral Entropy of Resting-State EEG,” Brain Topogr, vol. 28, no. 5, pp. 680–690, Sep. 2015, doi: 10.1007/s10548-015-0429-3.

[18] K. S. Lashley, Brain mechanisms and intelligence: A quantitative study of injuries to the brain. Chicago: University of Chicago Press, 1929. doi: 10.1037/10017-000.

[19] A. M. Bastos and J.-M. Schoffelen, “A Tutorial Review of Functional Connectivity Analysis Methods and Their Interpretational Pitfalls,” Front. Syst. Neurosci., vol. 9, Jan. 2016, doi: 10.3389/fnsys.2015.00175.

[20] J. Gonzalez-Astudillo, T. Cattai, G. Bassignana, M.-C. Corsi, and F. De Vico Fallani, “Network-based brain–computer interfaces: principles and applications,” J. Neural Eng., vol. 18, no. 1, p. 011001, Feb. 2021, doi: 10.1088/1741-2552/abc760.

[21] E. C. W. Van Straaten and C. J. Stam, “Structure out of chaos: Functional brain network analysis with EEG, MEG, and functional MRI,” European Neuropsychopharmacology, vol. 23, no. 1, pp. 7–18, Jan. 2013, doi: 10.1016/j.euroneuro.2012.10.010.

[22] D. S. Bassett and O. Sporns, “Network neuroscience,” Nat Neurosci, vol. 20, no. 3, pp. 353–364, Mar. 2017, doi: 10.1038/nn.4502.

[23] R. Zhang et al., “Efficient resting-state EEG network facilitates motor imagery performance,” J. Neural Eng., vol. 12, no. 6, p. 066024, Dec. 2015, doi: 10.1088/1741-2560/12/6/066024.

[24] F. Li et al., “Brain Network Reconfiguration During Motor Imagery Revealed by a Large-Scale Network Analysis of Scalp EEG,” Brain Topogr, vol. 32, no. 2, pp. 304–314, Mar. 2019, doi: 10.1007/s10548-018-0688-x.

[25] M. Lee, J.-G. Yoon, and S.-W. Lee, “Predicting Motor Imagery Performance From Resting-State EEG Using Dynamic Causal Modeling,” Front. Hum. Neurosci., vol. 14, p. 321, Aug. 2020, doi: 10.3389/fnhum.2020.00321.

[26] N. Leeuwis, S. Yoon, and M. Alimardani, “Functional Connectivity Analysis in Motor-Imagery Brain Computer Interfaces,” Front. Hum. Neurosci., vol. 15, p. 732946, Oct. 2021, doi: 10.3389/fnhum.2021.732946.

[27] M. E. Raichle, “The Brain’s Default Mode Network,” Annu. Rev. Neurosci., vol. 38, no. 1, pp. 433–447, Jul. 2015, doi: 10.1146/annurev-neuro-071013-014030.

[28] J. R. Andrews-Hanna, “The Brain’s Default Network and Its Adaptive Role in Internal Mentation,” Neuroscientist, vol. 18, no. 3, pp. 251–270, Jun. 2012, doi: 10.1177/1073858411403316.

[29] R. L. Buckner and L. M. DiNicola, “The brain’s default network: updated anatomy, physiology and evolving insights,” Nat Rev Neurosci, vol. 20, no. 10, pp. 593–608, Oct. 2019, doi: 10.1038/s41583-019-0212-7.

[30] G. L. Colclough, M. W. Woolrich, P. K. Tewarie, M. J. Brookes, A. J. Quinn, and S. M. Smith, “How reliable are MEG resting-state connectivity metrics?,” NeuroImage, vol. 138, pp. 284–293, Sep. 2016, doi: 10.1016/j.neuroimage.2016.05.070.

[31] H. Cho, M. Ahn, S. Ahn, M. Kwon, and S. C. Jun, “EEG datasets for motor imagery brain–computer interface,” GigaScience, vol. 6, no. 7, p. gix034, Jul. 2017, doi: 10.1093/gigascience/gix034.

[32] M.-H. Lee et al., “EEG dataset and OpenBMI toolbox for three BCI paradigms: an investigation into BCI illiteracy,” GigaScience, vol. 8, no. 5, p. giz002, May 2019, doi: 10.1093/gigascience/giz002.

[33] E. Combrisson and K. Jerbi, “Exceeding chance level by chance: The caveat of theoretical chance levels in brain signal classification and statistical assessment of decoding accuracy,” Journal of Neuroscience Methods, vol. 250, pp. 126–136, Jul. 2015, doi: 10.1016/j.jneumeth.2015.01.010.

[34] G. R. Müller-Putz, R. Scherer, C. Brunner, R. Leeb, and G. Pfurtscheller, “Better than random? A closer look on BCI results”.

[35] B. Schelter, M. Winterhalder, and J. Timmer, Eds., Handbook of Time Series Analysis: Recent Theoretical Developments and Applications, 1st ed. Wiley, 2006. doi: 10.1002/9783527609970.

[36] L. Barnett and A. K. Seth, “The MVGC multivariate Granger causality toolbox: A new approach to Granger-causal inference,” Journal of Neuroscience Methods, vol. 223, pp. 50–68, Feb. 2014, doi: 10.1016/j.jneumeth.2013.10.018.

[37] K. Hlavackovaschindler, M. Palus, M. Vejmelka, and J. Bhattacharya, “Causality detection based on information-theoretic approaches in time series analysis,” Physics Reports, vol. 441, no. 1, pp. 1–46, Mar. 2007, doi: 10.1016/j.physrep.2006.12.004.

[38] G. Nolte et al., “Robustly Estimating the Flow Direction of Information in Complex Physical Systems,” Phys. Rev. Lett., vol. 100, no. 23, p. 234101, Jun. 2008, doi: 10.1103/PhysRevLett.100.234101.

[39] L. Barnett, A. B. Barrett, and A. K. Seth, “Granger Causality and Transfer Entropy Are Equivalent for Gaussian Variables,” Phys. Rev. Lett., vol. 103, no. 23, p. 238701, Dec. 2009, doi: 10.1103/PhysRevLett.103.238701.

[40] R. Oostenveld, P. Fries, E. Maris, and J.-M. Schoffelen, “FieldTrip: Open Source Software for Advanced Analysis of MEG, EEG, and Invasive Electrophysiological Data,” Computational Intelligence and Neuroscience, vol. 2011, pp. 1–9, 2011, doi: 10.1155/2011/156869.

[41] P. L. Nunez et al., “EEG coherency,” Electroencephalography and Clinical Neurophysiology, vol. 103, no. 5, pp. 499–515, Nov. 1997, doi: 10.1016/S0013-4694(97)00066-7.

[42] S. B. Rutkove, “Introduction to Volume Conduction,” in The Clinical Neurophysiology Primer, A. S. Blum and S. B. Rutkove, Eds., Totowa, NJ: Humana Press, 2007, pp. 43–53. doi: 10.1007/978-1-59745-271-7_4.

[43] G. Nolte, O. Bai, L. Wheaton, Z. Mari, S. Vorbach, and M. Hallett, “Identifying true brain interaction from EEG data using the imaginary part of coherency,” Clinical Neurophysiology, vol. 115, no. 10, pp. 2292–2307, Oct. 2004, doi: 10.1016/j.clinph.2004.04.029.

[44] J.-P. Lachaux, E. Rodriguez, J. Martinerie, and F. J. Varela, “Measuring phase synchrony in brain signals,” Hum. Brain Mapp., vol. 8, no. 4, pp. 194–208, 1999, doi: 10.1002/(SICI)1097-0193(1999)8:4<194::AIDHBM4>3.0.CO;2-C.

[45] M. Rubinov and O. Sporns, “Complex network measures of brain connectivity: Uses and interpretations,” NeuroImage, vol. 52, no. 3, pp. 1059–1069, Sep. 2010, doi: 10.1016/j.neuroimage.2009.10.003.

[46] D. J. Watts and S. H. Strogatz, “Collective dynamics of ‘small-world’ networks,” Nature, vol. 393, no. 6684, pp. 440–442, Jun. 1998, doi: 10.1038/30918.

[47] V. Latora and M. Marchiori, “Efficient Behavior of Small-World Networks,” Phys. Rev. Lett., vol. 87, no. 19, p. 198701, Oct. 2001, doi: 10.1103/PhysRevLett.87.198701.

[48] K. Börner, S. Sanyal, and A. Vespignani, “Network science,” Annual Review Info Sci & Tec, vol. 41, no. 1, pp. 537–607, Jan. 2007, doi: 10.1002/aris.2007.1440410119.

[49] J. D. Power, B. L. Schlaggar, C. N. Lessov-Schlaggar, and S. E. Petersen, “Evidence for Hubs in Human Functional Brain Networks,” Neuron, vol. 79, no. 4, pp. 798–813, Aug. 2013, doi: 10.1016/j.neuron.2013.07.035.

[50] L. C. Freeman, “Centrality in social networks conceptual clarification,” Social Networks, vol. 1, no. 3, pp. 215–239, Jan. 1978, doi: 10.1016/0378-8733(78)90021-7.

[51] M. E. J. Newman, “Fast algorithm for detecting community structure in networks,” Phys. Rev. E, vol. 69, no. 6, p. 066133, Jun. 2004, doi: 10.1103/PhysRevE.69.066133.

[52] M. E. J. Newman, “Modularity and community structure in networks,” Proc. Natl. Acad. Sci. U.S.A., vol. 103, no. 23, pp. 8577–8582, Jun. 2006, doi: 10.1073/pnas.0601602103.

[53] E. A. Leicht and M. E. J. Newman, “Community Structure in Directed Networks,” Phys. Rev. Lett., vol. 100, no. 11, p. 118703, Mar. 2008, doi: 10.1103/PhysRevLett.100.118703.

[54] R. Guimerà and L. A. N. Amaral, “Cartography of complex networks: modules and universal roles,” J. Stat. Mech., vol. 2005, no. 02, p. P02001, Feb. 2005, doi: 10.1088/1742-5468/2005/02/P02001.

[55] C. H. Kasess, K. E. Stephan, A. Weissenbacher, L. Pezawas, E. Moser, and C. Windischberger, “Multisubject analyses with dynamic causal modeling,” NeuroImage, vol. 49, no. 4, pp. 3065–3074, Feb. 2010, doi: 10.1016/j.neuroimage.2009.11.037.

[56] C. H. Kasess, C. Windischberger, R. Cunnington, R. Lanzenberger, L. Pezawas, and E. Moser, “The suppressive influence of SMA on M1 in motor imagery revealed by fMRI and dynamic causal modeling,” NeuroImage, vol. 40, no. 2, pp. 828–837, Apr. 2008, doi: 10.1016/j.neuroimage.2007.11.040.

[57] M. Bönstrup, R. Schulz, J. Feldheim, F. C. Hummel, and C. Gerloff, “Dynamic causal modelling of EEG and fMRI to characterize network architectures in a simple motor task,” NeuroImage, vol. 124, pp. 498–508, Jan. 2016, doi: 10.1016/j.neuroimage.2015.08.052.

[58] J. P. Kuhtz-Buschbeck, C. Mahnkopf, C. Holzknecht, H. Siebner, S. Ulmer, and O. Jansen, “Effector-independent representations of simple and complex imagined finger movements: a combined fMRI and TMS study,” Eur J of Neuroscience, vol. 18, no. 12, pp. 3375–3387, Dec. 2003, doi: 10.1111/j.14609568.2003.03066.x.

[59] T. Zhang et al., “Structural and functional correlates of motor imagery BCI performance: Insights from the patterns of fronto-parietal attention network,” NeuroImage, vol. 134, pp. 475–485, Jul. 2016, doi: 10.1016/j.neuroimage.2016.04.030.

[60] Y. K. Kim, E. Park, A. Lee, C.-H. Im, and Y.-H. Kim, “Changes in network connectivity during motor imagery and execution,” PLoS ONE, vol. 13, no. 1, p. e0190715, Jan. 2018, doi: 10.1371/journal.pone.0190715.

